# Dopamine neurons gate the intersection of cocaine use, decision making, and impulsivity

**DOI:** 10.1101/2020.06.29.179069

**Authors:** Tristan Hynes, Kelly Hrelja, Brett Hathaway, Celine Hounjet, Chloe Chernoff, Sophie Ebsary, Graeme Betts, Brittney Russell, Lawrence Ma, Sukhbir Kaur, Catharine Winstanley

## Abstract

Gambling and substance use disorders are highly comorbid. Both clinical populations are impulsive and exhibit risky decision-making. Drug-associated cues have long been known to facilitate habitual drug-seeking, and the salient audiovisual cues embedded within modern gambling products may likewise encourage problem gambling. The dopamine neurons of the ventral tegmental area (VTA) are exquisitely sensitive to drugs of abuse, uncertain rewards, and reward-paired cues, and may therefore be the common neural substrate mediating synergistic features of both disorders. To test this hypothesis, we first gained specific inhibitory control over VTA dopamine neurons by transducing a floxed inhibitory DREADD (AAV5-hSyn-DIO-hM4D(Gi)-mCherry) in rats expressing Cre recombinase in tyrosine hydroxylase neurons. We then trained rats in our cued rat gambling task (crGT), inhibiting dopamine neurons throughout task acquisition and performance, before allowing them to self-administer cocaine in the same diurnal period as crGT sessions. The trajectories of addiction differ in women and men, and the dopamine system may differ functionally across the sexes, therefore we used male and female rats here. We found that inhibition of VTA dopamine neurons decreased cue-induced risky choice and reduced motor impulsivity in males, but surprisingly, enhanced risky decision making in females. Inhibiting VTA dopamine neurons also prevented cocaine-induced deficits in decision making in both sexes, but nevertheless drove all animals to consume more cocaine. These findings show that chronic dampening of dopamine signalling can have both protective and deleterious effects on addiction-relevant behaviours, depending on biological sex and dependent variable of interest.

## 1. Introduction

The choice to use illicit, addictive drugs such as cocaine is inherently risky, and often impulsive. It is therefore unsurprising that high levels of risk taking and impulsivity are associated with substance use disorder (SUD) (1–3). Risk taking and impulsivity are also strongly associated with gambling disorder (GD). Furthermore, GD is often comorbid with impulse control problems and chemical dependencies. Theories of both GD and SUD posit a critical role for cues (4, 5). Gamblers can respond to the allure of flashy casino light and sounds with a craving to gamble, similar to the way in which drug users react to the cues associated with their drug of choice (e.g., a syringe or crack pipe). Thus, cue responsivity appears to be an important link between these disorders. To push forward development of treatments for SUD and GD, understanding the precise neurobiological mechanisms through which cues promote addiction vulnerability is vitally important.

In order to probe the neurobiology that underlies the interplay between cues, GD, and SUD, we have combined our cued rat gambling task with operant cocaine self-administration. In the rat gambling task (rGT), rats learn to respond at an array of illuminated response apertures to earn sugary rewards. Similar to human gambling paradigms, the “high-risk, high-reward” options yield larger potential per-trial gains, but also greater and more frequent time-out penalties. The optimal strategy is therefore to favor options that deliver smaller but more certain rewards, and fewer, lesser punishments. The vast majority of rats employ an optimal strategy, as this maximizes the total amount of food they receive. Much like in real-world casinos, and in experimental gambling tasks for humans (6, 7), adding casino-inspired sound and light cues to wins in the rGT causes more rats to choose the risky options (8). The addition of these casino-like cues to the rGT also enhances the ability of volitionally self-administered cocaine to sway choice toward the risky options (9). Moreover, rats trained in the cued rGT appear more motivated to take cocaine in general. This evidence suggests that the presence of gambling-like cues may facilitate the interaction between risk taking and cocaine use. As such, the ability of reward-concurrent cues to promote cocaine use and problem gambling may be underpinned by similar neurobiological mechanisms, partly explaining why GD and SUD are highly comorbid.

Animals that learned the cued rGT (in which risk taking is more prevalent) also exhibited lower levels of tonic dopamine in the nucleus accumbens (NAc) at the end of training (9). Similarly, repeated administration of cocaine reduces tonic NAc dopamine levels, as well as phasic NAc dopamine responses to cues and cocaine (10–13). It therefore seems that repeated exposure to either heavily-cued probabilistic rewards or psychostimulant drugs, both of which acutely induce dopamine release, may lead to a down-regulation of both tonic and phasic dopaminergic activity. Cue- and cocaine-induced suppression of dopamine may lead to reward deficient states, and drive engagement in dopamine-evoking activities, like risk taking and cocaine use. Collectively, these data lead us to the following hypothesis: if the dopamine-evoking activity of cues could be reduced while rats learn the cued rGT, cue-induced risk taking could be attenuated. Similarly, if cocaine experience is promoting risk-taking through a similar mechanism, dampening dopamine during the cued rGT sessions that follow cocaine self-administration training should also prevent the concomitant increase in risky choice. To test our hypotheses, we gained inhibitory chemogenetic control over the dopamine neurons projecting from the VTA. These neurons are exquisitely responsive to reward-paired cues and probabilistic rewards, and should therefore be strongly activated during the cued rGT. If this reduction in task-evoked VTA dopamine neuron activity prevents compensatory reductions in dopamine release, the neuroadaptations leading to reward deficiency should be attenuated. Thus, we predicted that chemogenetic inhibition of VTA dopamine neurons during cued rGT acquisition should diminish risk taking at baseline. Similarly, dampening VTA dopamine response on task should also prevent cocaine experience from enhancing risky choice. Mesolimbic dopamine levels also tightly regulate motor impulsivity, as assessed by rats’ inability to wait for the response apertures to illuminate before responding (14, 15). We therefore anticipated that chemogenetic inhibition of VTA dopamine neurons would reduce such premature responding. Regarding predicted sex differences, females are thought to have more sensitive dopamine systems, but recent data suggest dopamine antagonists are less effective at modulating decision making on the rGT in female rats (16, 17). Drug addiction also takes a different trajectory in females (18, 19). We therefore predicted that the effect of suppressing VTA dopamine neuron activity may be greater in males than females.

## 2. Methods

### Subjects

Subjects were 32 male (transgene positive (TG+): n = 16; transgene negative (TG−): n = 16) and 32 female (TG+: n = 16; TG−: n = 16) transgenic rats, bred in house against a Long-Evans background (Charles River, St. Constant, QC), that expressed cre recombinase (Cre) in neurons containing tyrosine hydroxylase (Long-Evans-TG(TH-Cre)3.1Deis, RRRC #00659; Rat Resource and Research Centre, RRRC, Columbia, MO). Rats were pair- or trio-housed in a climate-controlled colony room on a reverse 12 hours light-dark cycle (lights off 08:00; temperature 21°C). All housing conditions and testing procedures were in accordance with the guidelines of the Canadian Council on Animal Care, and all protocols were approved by the Animal Care Committee of the University of British Columbia.

### 2.1 Experimental Timeline

Over the course of a 2-week period, rats underwent stereotaxic surgery for delivery of the viral vector. All rats recovered with their cage mates for 4-6 weeks, and two weeks prior to commencement of crGT training, were food restricted to 85% of their free feeding weight. crGT began 4-6 weeks after surgery. Thirty minutes prior to each pre-training and crGT session (which took place from 09:00 – 09:30), rats received IP injections of CNO (1.0 mg/kg). After 35 crGT sessions, rats were implanted with jugular vein catheters and randomly assigned to cocaine or saline self-administering groups. After allowing 1 week for recovery, rats again began daily crGT sessions, with CNO still administered prior. After 5 baseline sessions of crGT, rats began cocaine self-administration sessions, which took place from 17:00 – 19:00. crGT testing was conducted the following morning from 09:00 – 09:30. CNO was still administered to all rats prior to the crGT sessions. CNO has been shown to effectively modulate behaviour and activate DREADDs for 40-70 minutes (20, 21). As such, by spacing the CNO injections and self-administration sessions apart by 8.5 hours, we minimised the chances that CNO would be active during the cocaine/saline self-administration sessions. After 10 consecutive days of cocaine self-administration with concurrent crGT sessions, rats were euthanized and their brains were processed for immunohistochemistry (Figure 1).

**Fig. 1.**
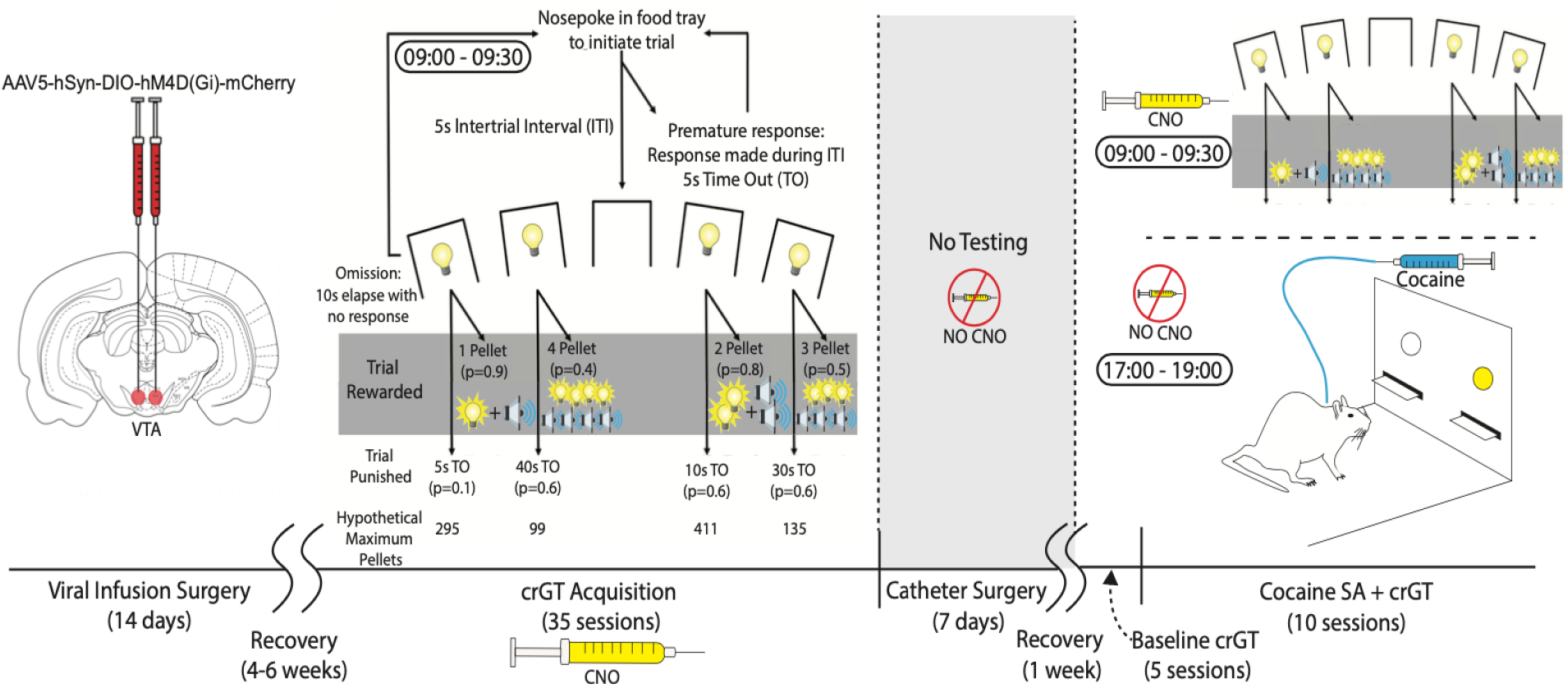
Experimental Timeline. The inhibitory DREADD AAV5-hSyn-DIO-hM4D(Gi)-mCherry was sterotaxically delivered into the bilateral VTA of female and male TH::Cre rats, which were then given 4-6 weeks to recover with their cage mates with ad libitum food. Rats were then food restricted and began daily crGT training, where CNO (1.0 mg/kg; i.p.) was delivered 30 minutes prior to the commencement of each session. After completing 35 sessions of crGT, rats were implanted with jugular vein catheters, singly housed, and allowed to recover in their home cages for one week. Rats then completed 5 baseline sessions of crGT with CNO on board, before entering into a phase of concurrent cocaine self-administration and crGT training. Each morning, CNO was administered 30 minutes prior to crGT training. Following each crGT session, rats were returned to their cages for 8 hours and, later that evening, placed in different operant boxes and allowed to self-administer intravenous cocaine for 2 hours (0.5 mg/kg/infusion; FR1). After 10 consecutive days of concurrent crGT and cocaine self-administration, rats were euthanized, and their brains were processed for immunohistochemistry.

### 2.2 Stereotaxic Surgery

We induced anesthesia with 5% isofluorane in oxygen (2 L/min flow rate) before securing rats in a stereotaxic frame and bilaterally infusing AAV5-hSyn-DIO-hM4D(Gi)-mCherry (0.5 uL/hemisphere; UNC Vector Core) into the VTA (AP: −5.5, ML:+/−0.6, DV:-8.2 from skull) at a rate of 0.10 uL/min using standard stereotaxic techniques. We left the injector tip in place for a further 5 minutes to allow for diffusion of the bolus. All rats received 5 mg/kg ketaprofen every 12 hours for 48 hours following surgery. Animals were then maintained on standard rat chow (14 g/day/male; 11 g/day/female), plus the sugar pellets earned in the task (~5 g per day). Water was available ad libitum at all times. Behavioural testing began at least one week following the start of food restriction.

### 2.3 Apparatus

The crGT and cocaine self-administration were each conducted in 16 separate banks of operant conditioning chambers (30.5 × 24 × 21 cm; Med Associates, St. Albans, VT, USA), located in separate rooms. Each chamber was enclosed within a ventilated sound-attenuating cabinet (Med Associates Inc, VT). The operant boxes were equipped with a fan to provide ventilation and to mask extraneous noise. Set in the curved wall of each box was an array of five holes. Each nose-poke unit was equipped with an infrared detector and a yellow light-emitting diode stimulus light. Sucrose pellets (45 mg, Formula P; Bio-Serv) could be delivered at the opposite wall via a dispenser. The cocaine self-administration chambers were identical to the crGT chambers with the exception of an additional cantilevered drug deliver arm and vascular access tether system plus externally mounted variable rate syringe pump. Online control of the apparatus and data collection was performed using code written by CAW in MEDPC (Med Associates) running on standard IBM-compatible computers.

### 2.4 The Cued Rat Gambling Task (crGT)

All rats were trained on the cued rat gambling task (crGT), as described previously 8. Animals were initially habituated to the operant chambers over the course of two 30 min exposures during which sucrose pellets were placed in each of the apertures and animals were allowed to explore the apparatus. Animals were then trained on a variant of the five-choice serial-reaction time task (5-CSRTT) in which one of the five nose-poke apertures was illuminated for 10 s and a nose-poke response was rewarded with a single sucrose pellet delivered to the food magazine. The aperture in which the stimulus light was illuminated varied across trials. Each session consisted of 100 trials and lasted 30 min. Animals were trained on this task until responding reached 80% accuracy and 20% omissions. Once this training was complete, rats then performed a forced-choice variant of the crGT. This training procedure ensured equal exposure to the different reinforcement contingencies associated with each aperture. During each 30-minute crGT session, rats sampled between four response holes, each of which was associated with distinct magnitudes and probabilities of sucrose pellet rewards or time-out punishments (see Figure 1). The optimal approach in the crGT is to favour options which deliver smaller per-trial gains but lower time-out penalties; consistent choice of the smaller reward options was advantageous due to more frequent rewards, but also less frequent and shorter time-outs, with the two-pellet choice (P2) resulting in the most reward earned per unit time. As detailed in our previous publications, most rats adopt this optimal strategy in the absence of reward-concurrent audiovisual cues (22). A 2-s audiovisual cue was presented concurrent with reward delivery on each option. The cue increased in complexity with the size of the reward: P1 win: P1 hole flashes for 2s at 2.5 Hz, monotone; P2 win: P2 hole flashes for 2s at 2.5 Hz, tone changes pitch once after 1s; P3 win: P3 hole flashes at 5 Hz for 1s, followed by flashing of the two neighboring holes in one of two patterns chosen at random, traylight flashes concurrently at 0.2 Hz for 2s, three different tones used, changing pitch every 0.1 s, in one of two patterns chosen at random; P4 win: P4 hole flashes at 5 Hz for 1s, followed by flashing of all five holes in one of four patterns at random, traylight flashes concurrently at 0.2 Hz for 2s, six different tones used, changing pitch every 0.1 s, in one of four patterns chosen at random.

Responses made during the ITI were recorded as premature responses, a measure of impulsive action, which resulted in the illumination of the house light and a 5 s time-out penalty after which a new trial could be initiated. If a response was not made into one of the 4 holes during the 10 s stimulus presentation, the trial would be registered as an omission, after which point another trial would begin. Animals received one training session per day, 5-7 days per week, until statistically stable, asymptotic levels of performance were observed. That is, animals were considered to have learned the task if no within-subjects effect of session was observed over 5 consecutive days of training (35 sessions).

### 2.5 Jugular catheter implantation

Once stable behaviour had been established on the crGT, animals were implanted with a intravenous catheter to enable drug self-administration. Specifically, we aseptically implanted catheters constructed of Silastic silicone tubing (Dow Corning via VWR International, Edmonton, AB, Canada), attached to back-mounted cannulae (Plastics One, Roanoke, VA, USA), into the right jugular vein as per our previous published methods (9). We passed the catheters through the skin subcutaneously and externalized the cannulae between the scapulae. Following surgery, the catheter was filled with a heparin glycerol lock solution (SAI Infusion Technologies, Illinois, USA) and the animals were left to recover for 5-7 days, after which crGT testing resumed and cocaine self-administration began. The day prior to the first cocaine self-administration session, catheter patency was tested by delivering 0.1 mL of 10% ketamine HCL (Medisca Pharmaceuticals, British Columbia, Canada). All rats exhibited an immediate loss of muscle tone and sedation, resulting in no rats being excluded on the criterion of catheter patency.

### 2.6 crGT-concurrent cocaine self-administration

Animals were trained to lever press for cocaine hydrochloride (0.50 mg/kg/infusion, dose calculated as the salt and dissolved in sterile 0.9% saline; Medisca Pharmaceuticals, British Columbia, Canada) or saline vehicle over 10 daily 2- hr sessions (23). At the start of each self-administration session, two free infusions of solution were given to fill catheters and indicate drug was available. Rats were presented with two levers, one active and one inactive, with an illuminated cue-light situated over the active lever. Using a fixed ratio (FR1) schedule, responses on the active lever would result in a single 4.5 s infusion in concert with the cue light flashing (50 Hz) and a novel 20 kHz tone (this tone was not used in the crGT). Following the infusion, animals underwent a 10 s time-out during which drug would not be delivered, the cue light and tone were not presented, but levers would remain extended and responses monitored. Responses on the active lever during infusions and timeouts were recorded and interpreted as preliminary cocaine “seeking” behaviors. Inactive lever presses, while monitored, had no programmed consequences. Animals were limited to 30 infusions per hour to prevent overdose.

### 2.7 Histology

In order to visualize DREADD expression in VTA DA neurons, we double-labelled 35um sections coronal sections with primary antibodies against mCherry (Cat#ab205402; Abcam; Toronto, ON, Canada; 1:700 for 48 hours) and tyrosine hydroxylase (Cat#AB152; Millipore Sigma; Oakville, ON, Canada; 1:100 for 24 hours) for 24 hours. Sections were then washed in PBS and incubated with secondary antibodies conjugated to Alexa Fluor ® 488 (Cat#A-21103) and Alexa Fluor ® 633 (Cat#A-11034) (Thermo Fischer Scientific; Burnaby, BC, Canada; 1:500 for both). Sections were then cover-slipped under HARLECO ® Krystalon™ mounting medium (Thermo Fischer Scientific; Burnaby, BC, Canada) and visualized using a SP8 WLL confocal microscope (Leica Microsystems, Germany). For quantification of DREADDs expression, we adopted a method similar to previous published methods (17, 24, 25). We defined the anterior and posterior bounds of DREADD expression (i.e., < 1 mCherry+ cell) in coronal sections and counted cells from 1-3 sections within these bounds immediately anterior or posterior to the DREADD infusion site. In both hemispheres, we quantified the number of neurons somatically co-expressing mCherry and TH, as well as those expressing mCherry alone. We averaged the number of cell bodies identified across hemispheres and sections for individual rats. We confirmed, but did not quantify, terminal expression in the nucleus accumbens and medial prefrontal cortex.

### 2.8 Statistical Analysis

We analyzed the following crGT variables per session: score [(*P*1 + *P*2) − (*P*3 + *P*4)], percent choice of each option (number of PX chosen/ total number of choices x 100), percentage of premature responses (number of premature responses/ total number of trials initiated x 100), sum of omitted responses, sum of trials completed, and average latencies to choose an option and collect reward. For cocaine self-administration analyzed number of infusions of achieved, number of responses on the active lever, and number of responses on the inactive lever. We used SPSS Statistics 25.0 (IBM; Chicago, IL, USA) to conduct all analyses. We performed repeated measures ANOVA over 35 sessions of task performance with sex and transgene status as between-subjects factors. We followed up significant within-subjects omnibus outcomes by comparing rates of task acquisition via linear contrasts. Significant between-subjects effects were up with independent samples t-tests. Total cocaine consumed and active responses made was calculated as the sum of all infusions achieved or responses made over the 10 day testing period. Because clinical and pre-clinical research suggests that impulsivity and decision making share a common latent construct (3, 26, 27), we wished to control for the relationship in some of the present analyses. In this particular group of rats, premature responding accounted for 7% of the variance in decision making score (*r*_59_ = −0.256, *p* = 0.025). For all ANCOVA analyses, we set the covariate as the average score over the last 5 days of task acquisition.

## 3. Results

### 3.1 Histology

Figures 2A-C provide evidence of DREADD expression in putative dopamine neurons throughout the VTA, with minimal expression in the substantia nigra (SNc). We detected some DREADD-expressing cells in the medial aspects of the SNc in 7 males and 4 females; in these instances, a maximum of 7 cells/section were observed. Figures 2D-F show that the majority of DREADD-expressing neurons co-expressed TH [% co-labelled cells = 92.62 +/− 4.42 (SEM)], indicating that DREADD expression was highly selective to dopamine neurons. Four TG−females and 6 TG− males displayed DREADD expression in midbrain TH+ neurons; we detected a maximum of 6 cells/section in these cases. Female and male TG+ rats did not differ in the number of VTA DREADD expressing cells (Figure 2 G&H; sex: *t*_27_ = 0.724, *p* = 0.476). We excluded from all analyses rats that expressed mCherry unilaterally (n = 1 TG+ male) or not at all (n = 3 TG+ females). One TG− female died post-operatively. We additionally imaged two of the main terminal fields of the dopaminergic projections from the VTA. Figure S1 shows the extent of putative dopaminergic innervation to the striatum and medial prefrontal cortex (mPFC). These terminal fields also robustly expressed DREADDs, suggesting that the effects we observe here may be mediate by activity of dopamine in both the striatum and the mPFC.

**Fig. 2.**
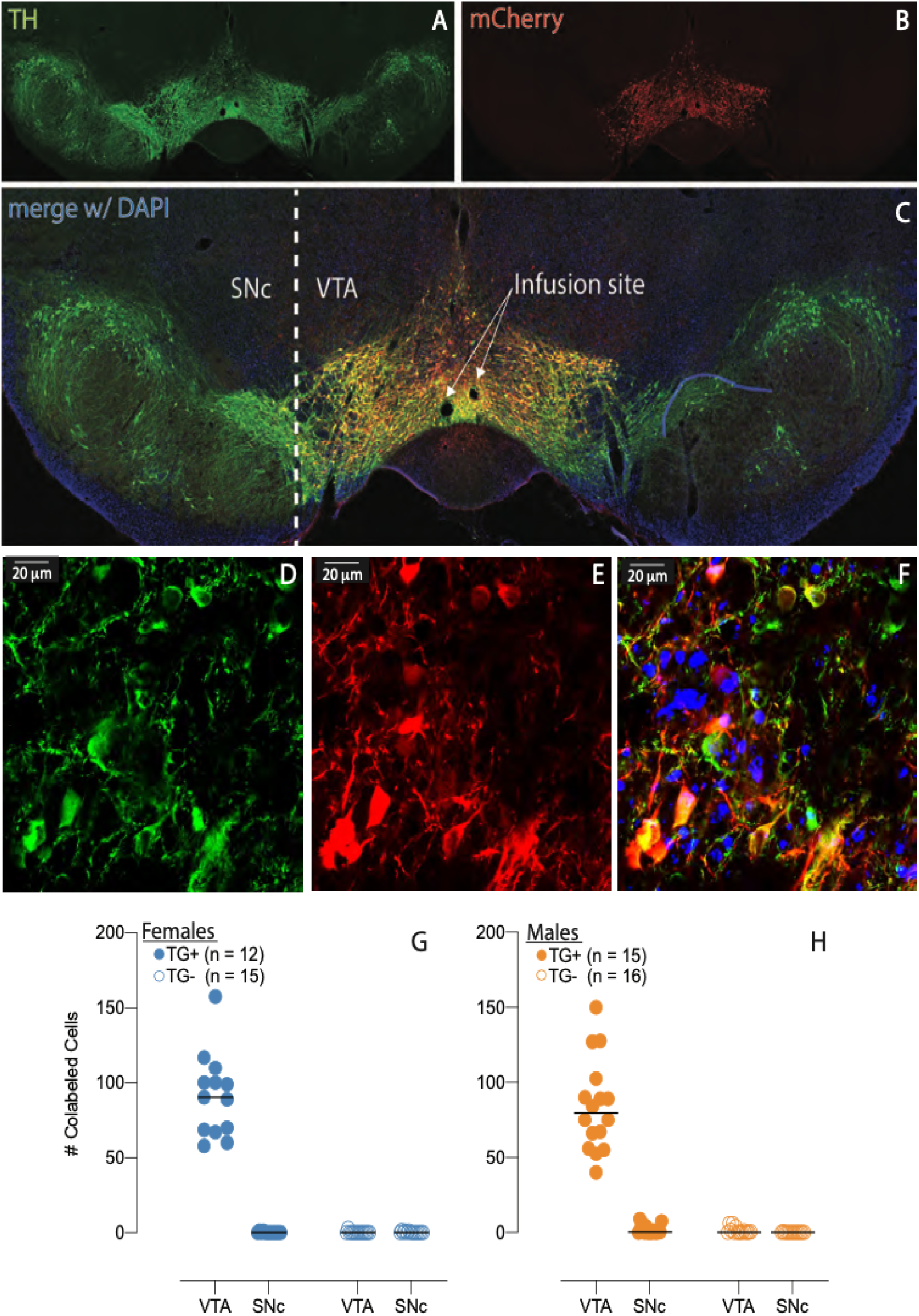
DREADD expression in VTA dopamine neurons. (A) 10X photomicrograph showing in green putative dopaminergic [i.e., tyrosine hydroxylase (TH) positive] cell bodies of the VTA. (B) Shown in red are mCherry-tagged neurons, indicating DREADD expression. (C) Shows both channels merged with DAPI. (D-F) 64X photopicrograph showing robust colabelling with TH and mCherry in VTA cell bodies and arbours. (G) In female and (H) male rats, the majority DREADD expressing dopaminergic neurons were found in the VTA. Some mCherry-tagged cells were observed in the medial aspects of the SNc.

### 3.2 Decision making profile

Males and females did not initially (i.e., in first 5 sessions) differ in decision making score (session x sex: *F*_4,220_ = 0.482, *p* = 0.749; session x sex X TG: *F*_4,220_ = 0.647, *p* = 0.579), yet in light of our recent report that females and males respond differently to manipulations of the dopamine system in the crGT, we analyzed the decision making profile of males and females separately (17). We first analyzed decision making score, which is a composite metric used to quantify the preference for risky options. As briefly described in the methods section, decision making score is calculated as the difference between the percent of trials on which an animal makes an optimal choice (i.e., chooses P1 or P2) vs. a risky choice (i.e., P3 or P4). A score of 100 indicates a perfectly optimal choice profile, while a score of −100 indicates a perfectly risky one. In support of our initial hypothesis, we demonstrated that chemogenetic inhibition of VTA dopamine neurons during acquisition of the crGT results in more optimal decision making, at least in males (Figure 3B: linear session x TG: *F*_1,29_ = 4.664, *p* = 0.039). Specifically, linear regression analysis revealed that TG− males (*r* = −0.861 points/day) underwent a more rapid decline in overall decision making score than TG+ males (*r* = −0.6524 points/day; comparison of slopes: *F*_1,66_ = 294.5, *p* < 0.01). However, when rates of premature responding were covaried in the score analysis for males, the omnibus ANOVA gave a non-significant outcome (linear session X TG: *F*_1,28_ = 0.538, *p* = 0.469).

**Fig. 3.**
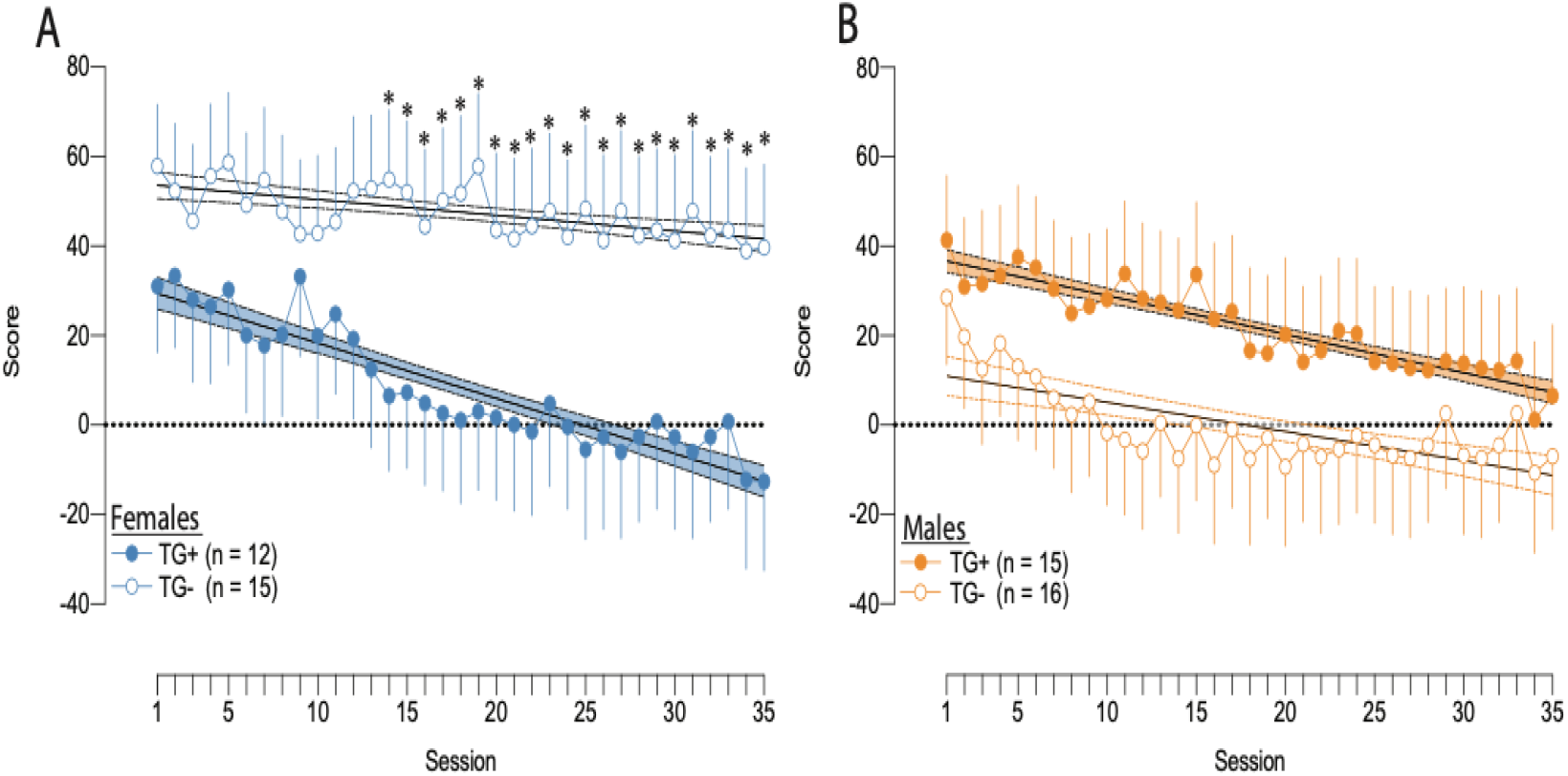
Longitudinal decision making score in female and male rats during crGT training. (A) TG+ females showed a steeper longitudinal decline in decision making ability than TG− females, following chronic administration of CNO prior to each crGT training session. (B) Male TG+ rats showed a showed a significantly slower decline in decision making ability than TG− rats, but did not significantly differ in absolute score, following acquisition. Data points represent group means +/− 1SEM. Linear regression line is shown intersecting data points and is bound by dashed lines, marking the 95% CIs. “*” indicates sessions on which independent samples t-tests showed TG+ and TG− rats to be significantly different.

In stark contrast to the males, TG+ females developed a significantly lower score over time than TG− females, indicative of greater risky choice (Figure 3A, session X TG: *F*_34,884_ = 1.728, *p* = 0.006). Using linear regression analyses to evaluate the change in score over session, we confirmed that TG+ females showed a more severe decline (*r* = −1.235 points/day) in overall decision making score than TG−females (*r* = −0.352 points/day) (comparison of slopes: *F*_1,66_ = 59.13, *p* < 0.001).Covarying premature response rate did not nullify the within-subjects effect in females, but did result in the detection of a between-subjects difference (*F*_1,25_ = 4.478, *p* = 0.044), which first emerged on session 14 and remained significant for the remainder of testing (TG− vs. TG+ – s1 to s13: all *ts* < 0.700, all *ps* > 0.491; s14: *t*26 = 2.115, *p* = 0.044; s35: *t*_26_ = 1.926, *p* = 0.033). Latent emergence of this effect is consistent with how the deleterious effect of cues on decision making only emerges over repeated testing sessions (28), suggesting that our present manipulation has enhanced the ability of the cues to affect decision making.

We also found further evidence of riskier decision making in females when activity of dopaminergic VTA neurons was dampened; TG+ females displayed a reduced preference for P2 (Figure 4C; session X transgene: *F*_34,850_ = 1.924, *p* = 0.001) and an increased preference for P4 (Figure 4H; session X TG: *F*_34,850_ = 2.891, *p* < 0.001; *TG* : *F*_1,25_ = 4.257, *p* = 0.050). These observations were further substantiated by subsequent linear regression analyses. Whereas TG− females showed an expected increased preference for the optimal choice P2 over time (*r* = 0.300), TG+ females exhibited a longitudinal decrease (*r* = −0.242) (*F*_1,66_ = 32.89, *p* < 0.001). TG+ females also exhibited a rapid increase in the preference for P4 over time (*r* = 0.337), whereas TG− females progressively decreased choice of this option (*r* = −0.096; comparison of slopes: *F*_1,66_ = 101.00, *p* < 0.0001). In summary, these results suggest that chemogenetically inhibiting dopaminergic neurons within the VTA caused severe deficits in decision making in females, and marginal improvements in males.

**Fig. 4.**
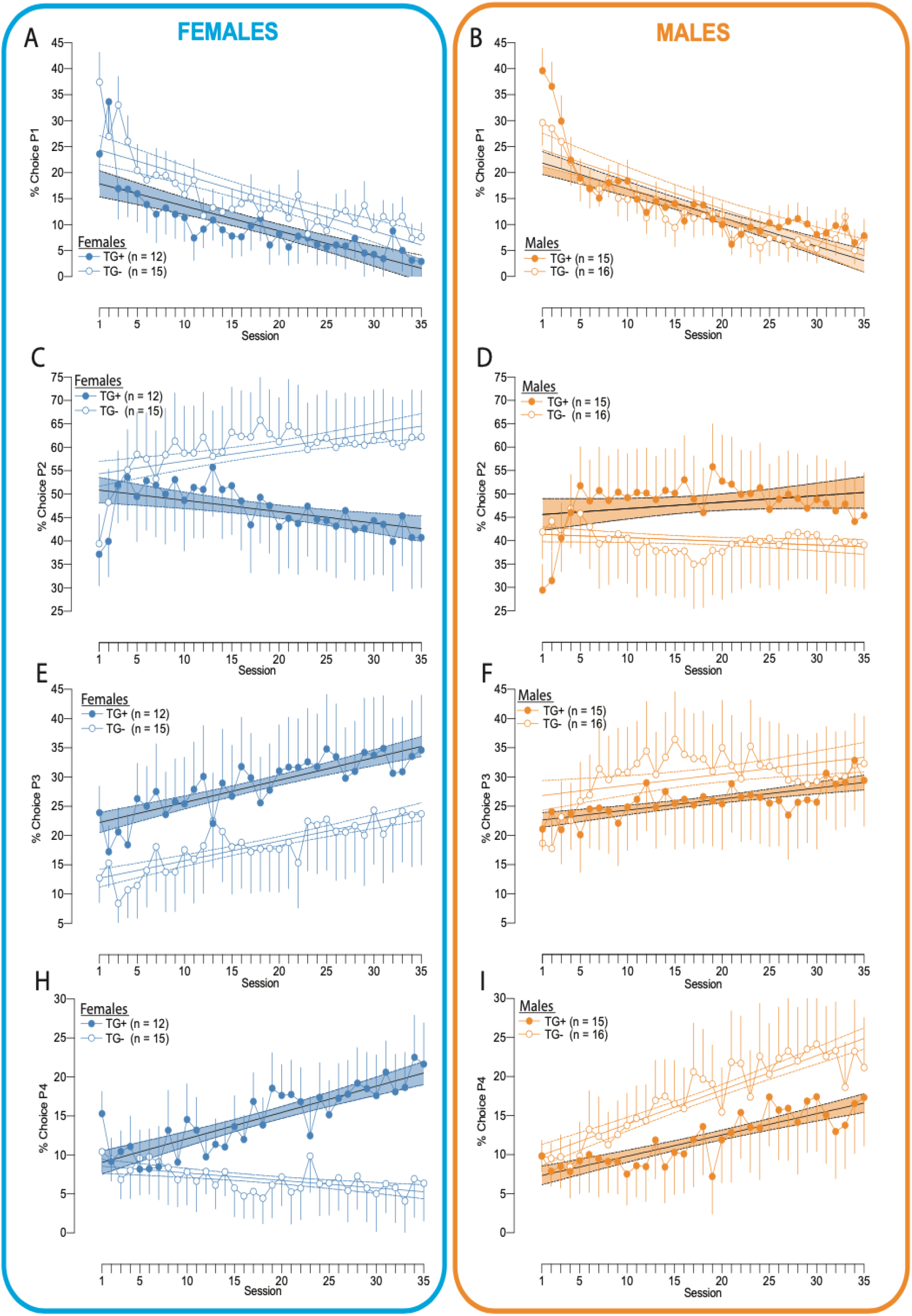
Longitudinal choice profile in female and male rats during crGT training. (A) TG+ females did not differ from TG− females in the propensity to choose the option P1. (B) TG+ and TG− males did not differ in the choice of P1. (C) TG− females acquired a preference for the optimal option P2 more rapidly than TG+ females. (D) TG+ males acquired a preference for the optimal choice, P2, more rapidly than TG− males. (E) TG+ and TG− females did not differ in the preference for P3. (F) TG+ and TG− males did not differ in their preference for P3 or (I) P4. (H) TG+ females acquired a preference for the disadvantageous option P4 more rapidly than their TG− counterparts. Data points represent group means +/− 1SEM. Linear regression line is shown intersecting data points and is bound by dashed lines, marking the 95% CIs.

### 3.3 Motor impulsivity

The omnibus ANOVA analysing premature responding revealed a between-subjects effects of transgene status (*F*_1,55_ = 10.356, *p* = 0.002) and of sex (*F*_1,55_ = 12.257, *p* = 0.001), again leading us to analyze females and males separately. Concordant with our hypotheses, TG+ males showed reduced premature responding throughout testing (Figure 5B; TG: *F*_1,29_ = 9.581, *p* = 0.004); these differences emerged on the 5^*th*^ session of crGT training and continued throughout (TG− vs. TG+ – s1-s4 : all *ts* < 0.967, all *ps* > 0.171; s5: *t*29 = 2.218, *p* = 0.018; s35: *t*_29_ = 0.699, *p* = 0.050). Transgene status did not impact premature responding in females (Figure 5A; *F*_1,26_ = 1.803, *p* = 0.191). The main effect of sex reported above appears to be driven by TG− males, perhaps suggesting baseline differences in impulsivity across the sexes, although this was not seen in a previous investigation (17).

**Fig. 5.**
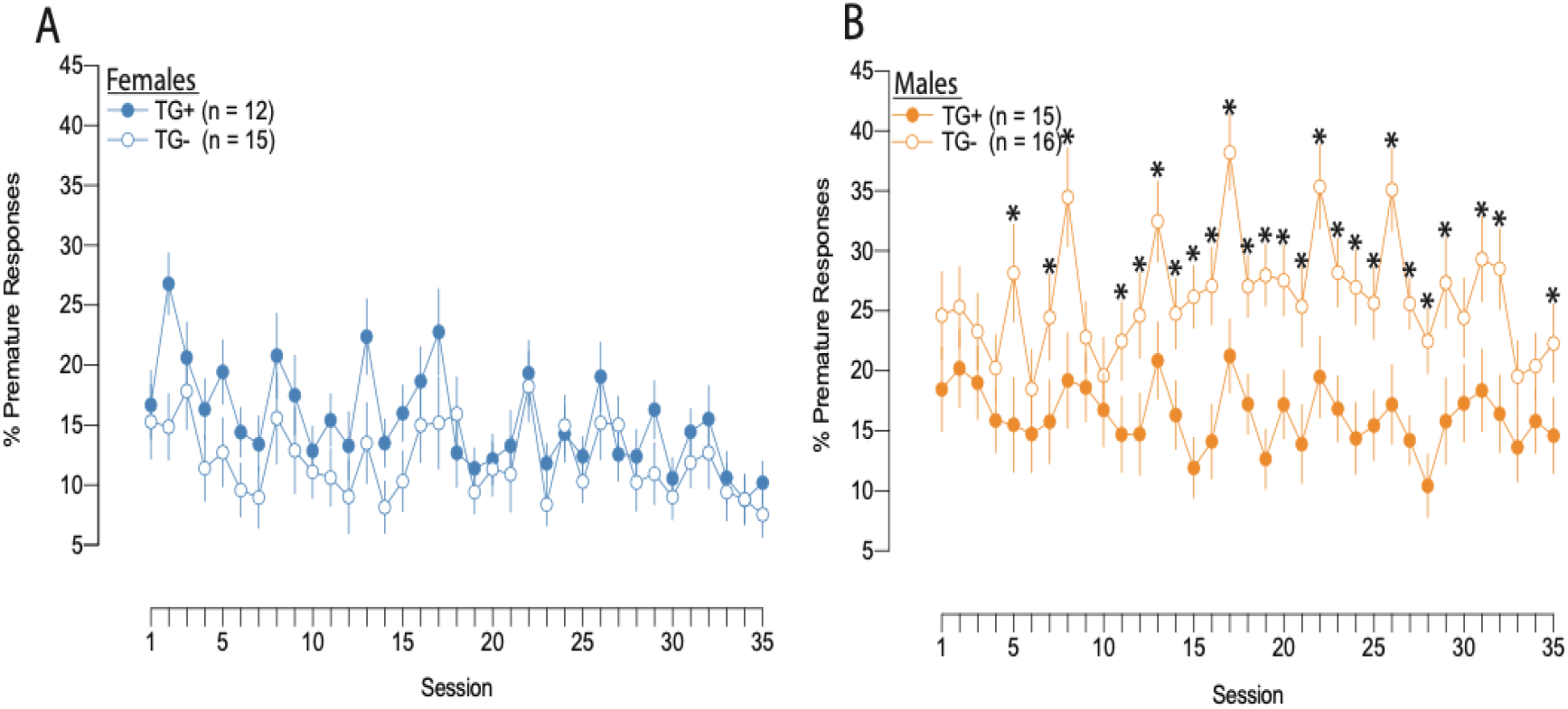
Longitudinal impulsivity in female and male rats during crGT training. (A) TG+ and TG− females did not differ from one another in the rate of premature responding, following chronic administration of CNO prior to each crGT training session. (B) Beginning on session 5 and lasting the duration of training, TG+ rats made impulsive responses than TG− rats following chronic CNO administration prior to each rGT training session. Data points represent group means +/− 1SEM. “*” indicates sessions on which independent samples t-tests showed TG+ and TG− rats to be significantly different.

### 3.4 Additional variables

Although females were generally slower to make a choice than males, TG+ rats of both sexes were slower to choose an option (Figure S2; all rats: TG: *F*_1,55_ = 11.268, *p* = 0.001), sex: (*F*_1,55_ = 12.218, *p* = 0.001); TG, females: TG: *F*_1,26_ = 4.246, *p* = 0.049; males: *F*_1,29_ = 7.298, *p* = 0.011). TG+ animals of both sexes also made more omissions, although this only reached trend-level significance in males (Figure S3, all rats: TG: (*F*_1,55_ = *S*3.101, *p* = 0.001; sex: *F*_1,55_ = 9.240, *p* = 0.004); TG, -females: (*F*_1,26_ = 8.966, -males: *F*_1,29_ = 3.832, *p* = 0.060). These data may suggest that inhibiting VTA dopamine neurons lead to a general decrease in response vigour. However, TG+ male rats were only marginally slower to collect reward than TG− animals, and female TG+ rats were even faster in this regard than their TG− counterparts (Figure S4; all rats: sex x TG: *F*_1,55_ = 6.660, *p* = 0.003; TG, -females: *F*_1,26_ = 3.119, *p* = 0.089, -males: *F*_1,29_ = 4.010, *p* = 0.055). Although female TG+ animals completed fewer trials, this likely results from the longer average trial length resulting from the more frequent, lengthy time-out penalties incurred through persistent choice of P4, rather than a drop in motor output (Figure S5; all rats: session x TG x Sex: *F*_34,1870_ = 1.475, *p* = 0.038; TG, -females: *F*_1,26_ = 6.356, *p* = 0.018, -males: *F*_1,29_ = 0.016, *p* = 0.899).

### 3.5 Cocaine self-administration

As with the other analyses, we looked at females and males separately on measures of cocaine self-administration. With respect to infusions, we detected a main effect of drug in females (Figures 6A&B; drug: *F*_1,25_ = 8.016, *p* = 0.009) and in males (Figures 6C&D; *F*_1,28_ = 12.441, *p* = 0.001). Follow up linear contrasts showed that in females, consumption of cocaine (Figure 6A; *F*_1,18_ = 21.422, *p* < 0.000) but not saline (Figure 6B; *F*_1,7_ = 3.734, *p* = 0.095) increased over time. Cocaine similarly acted as a potent operant reinforcer in males (cocaine: *F*_1,21_ = 15.514, *p* = 0.001; saline: *F*_1,7_ = 0.997, *p* = 0.351) (See Figures 6C&D). Although CNO was no longer active during the self-administration sessions, TG+ females (Figure 6A; session X TG: *F*_1,17_ = 5.775, *p* = 0.028) and males (Figure 6C; session X TG: *F*_1,20_ = 4.860, *p* = 0.039) self-administered more cocaine than TG− rats. For total responses upon the active lever, neither females (Figure 6F; linear session: *F*_1,6_ = 0.258, *p* = 0.629) nor males (Figure 6H; linear session: *F*_1,6_ = 0.014, *p* = 0.909) found saline reinforcing. In contrast, cocaine motivated operant responding in both sexes (Figures 6E&G; linear session females: *F*_1,17_ = 21.734, *p* < 0.000; males: *F*_1,20_ = 10.744, *p* = 0.004). TG status did not significantly impact the number of active responses in females or males (all *Fs* < 3.143, all *ps* > 0.098). As shown in Figures 6I-L, responding on the inactive lever did not differ between cocaine and saline self-administering groups or as a function of transgene status (session × drug females: *F*_9,234_ = 0.772, *p* = 0.643; session X drug X TG females: *F*_9,234_ = 1.356, *p* = 0.209; session x drug males: *F*_9,207_ = 0.612, *p* = 0.786; session X drug X TG: *F*_9,207_ = 0.819, *p* = 0.600).

**Fig. 6.**
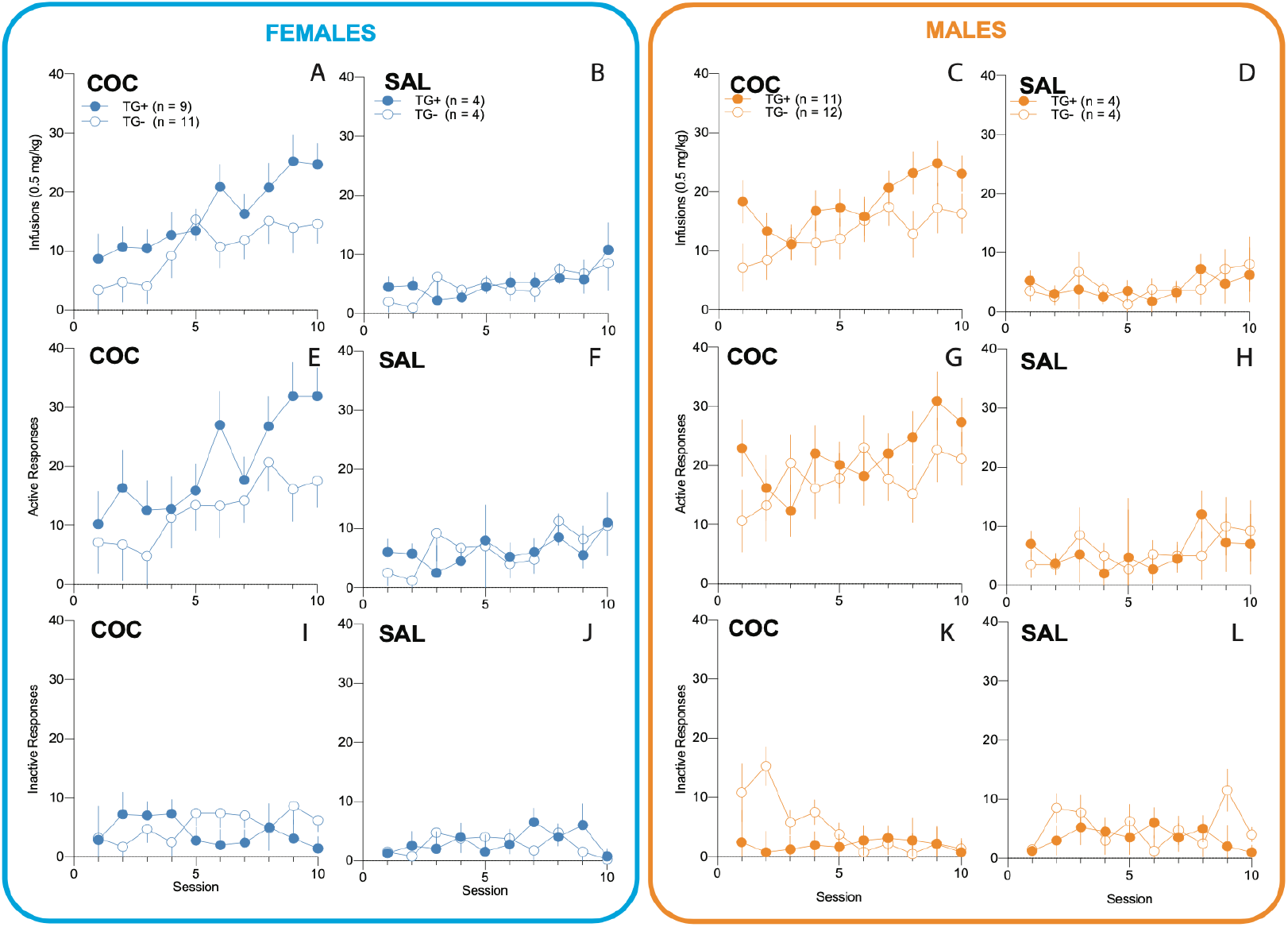
Cocaine self-administration profiles of female and male rats. (A) TG+ females took more infusions of cocaine than TG – females during the 2hr session. (B) Saline did not act as a reinforcer in TG+ or TG− females. (C) TG+ males took more infusions of cocaine than TG− males. (D) Saline did not act as a reinforcer in TG+ or TG− males. (E) TG+ and TG− females did not differ in the number of active responses made for cocaine or (F) saline. (G) TG+ and TG− females did not differ in the number of active responses made for cocaine or (H) saline. (I-L) Responding upon the inactive lever was not reinforcing for males or females, regardless of transgene status or whether saline or cocaine was self-administered.

**Fig. 7.**
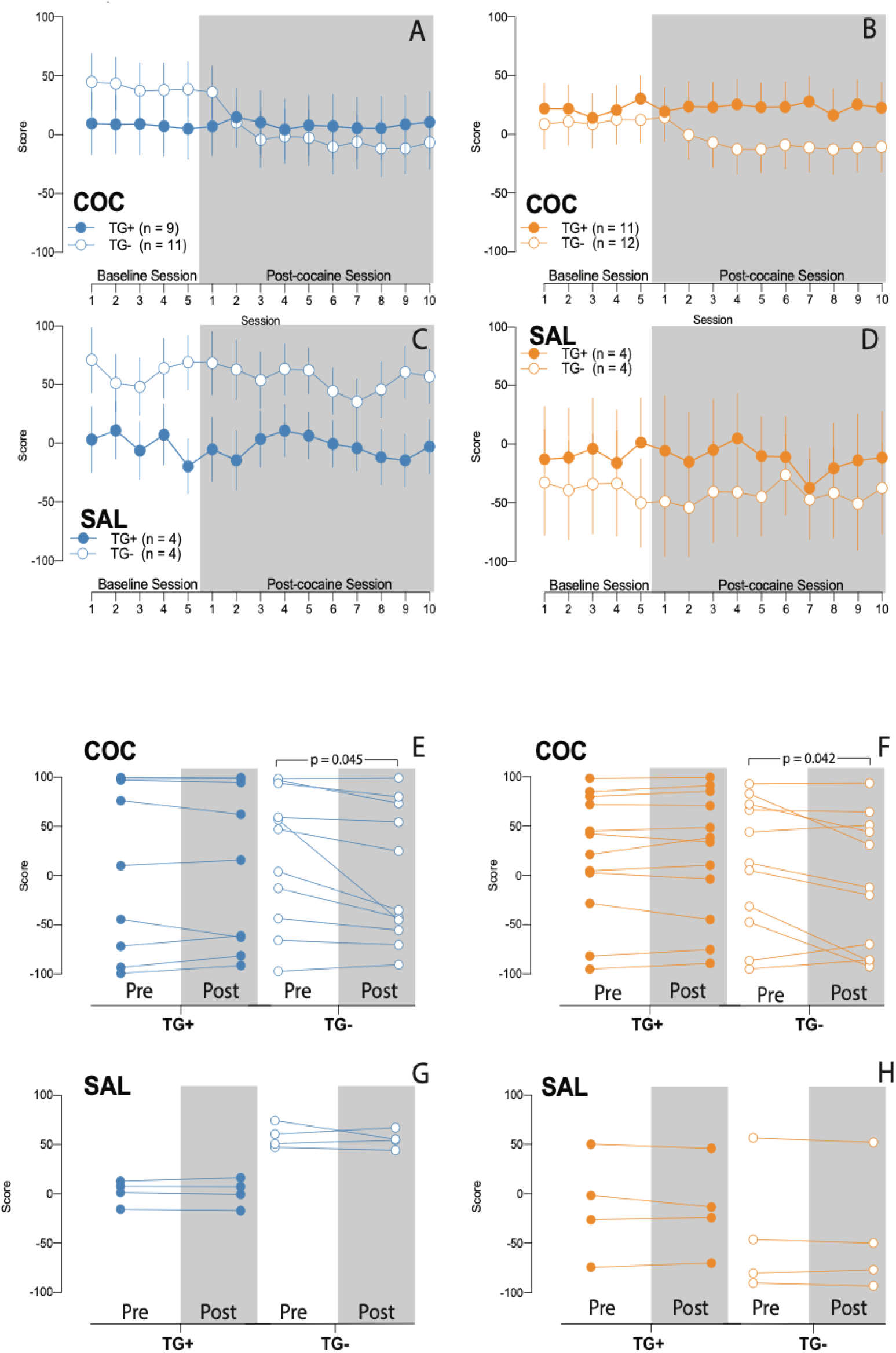
Effect of concomitant cocaine or saline self-administration on decision making. (A) TG− females and (B) TG− males underwent the expected reduction in decision making score following cocaine self-administration. (C) Females and (D) males were unaffected by saline self-administration had no effect on decision making score, regardless of transgene status. Direct comparisons of average score between pre- and post-cocaine substantiate the conclusion that (E) female and (F) male TG+ rats are resistant to cocaine-induced deficits in decision making. (G-H) As expected, saline self-administration had no effect on decision making score. Individual points with connecting lines represent the pre- and post-self-administration scores of individual animals. p-values indicate the outcome of 2-tailed paired-samples t-tests.

### 3.6 Effects of chronic CNO-mediated inhibition of VTA dopamine neurons on the ability of cocaine self-administration to drive risky choice

We have previously shown that 10 days of cocaine self-administration in the afternoons/evenings results in increased risky decision making on the crGT in males, as indicated by a decrease in score during the concomitant morning sessions. Although we observed the expected drop in score in TG− animals, this was not evident in TG+ rats, regardless of sex. Consistent with our approach throughout the manuscript, here we analyzed males and females separately. In females we detected the expected decrease in TG− rats, while TG+ rats were protected from the effect (linear session X transgene: *F*_1,18_ = 5.873, *p* = 0.026). We observed that TG+ males were likewise unaffected by cocaine, compared to the TG− males which made more risky choices (*F*_1,20_ = 10.275, *p* = 0.004). Saline self-administering rats exhibited no changes in decision making, regardless of transgene status (all *Fs* < 1.054, all *ps* > 0.410 We then collapsed data across sex and compared scores averaged over pre-cocaine (5 sessions) and post-cocaine (10 sessions) epochs of crGT performance, and confirmed a decrease in decision making score in female (*t*_10_ = 2.296, *p* = 0.045) and male (*t*_9_ = 2.368, *p* = 0.042) TG− rats but not TG+ rats (females: *t*_8_ = 0.206, *p* = 0.842; males: *t*_11_ = 0.751, *p* = 0.468). Saline self-administration did not alter decision making on the crGT (score: session X TG: *F*_14,140_ = 1.213, *p* = 0.273). We furthermore asked whether the amount of cocaine consumed was related to the magnitude of decline in decision making score, by correlating the total number of infusions achieved with the overall change in score at each level of sex and transgene status. None of the correlations reached statistical significance (all p’s < 0.106).

### 3.7 Effects of chronic CNO-mediated inhibition of VTA dopamine neurons on the ability of cocaine self-administration to decrease motor impulsivity

We and others have also observed that concurrent cocaine self-administration can reduce premature responding in male rats, consistent with theories surrounding the efficacy of psychostimulants in the treatment of impulse control disorders (29, 30). We conducted paired-samples t-tests, comparing the pre- and post-cocaine crGT training epochs at each level of sex, transgene status. As expected from our previous findings, we showed that TG− male rats exhibited reductions in motor impulsivity following cocaine self-administration (*t*_10_ = 2.528, *p* = 0.030), while TG+ males did not (*t*_11_ = 1.932, *p* = 0.079 Regardless of transgene status, females showed no reduction in premature responding (TG+: *t*_8_ = 0.138, *p* = 0.893; TG−: *t*_10_ = 2.032, *p* = 0.070). Saline self-administration had no impact on premature responding (all *ts* < 1.372, all *ps* > 0.264 (see Figure 6S).

### 3.8 Effects of chronic CNO-mediated inhibition of VTA dopamine neurons on the ability of cocaine self-administration to modulate other crGT variables

Following cocaine self-administration, TG+ female rats showed reductions in the number of trials completed (Figure S7A; session X TG: *F*_14,238_ = 2.424, *p* = 0.003; TG−: *t*_10_ = 3.104, *p* = 0.011; TG+: *t*_8_ = −0.826, *p* = 0.433), whereas males showed no difference (session X TG: *F*_14,294_ = 0.661, *p* = 0.812). Saline self-administration had no effect on trials competed in females (*F*_14,85_ = 0.994, *p* = 0.812) nor males (*F*_14,84_ = 0.797, *p* = 0.669) (see Figure S7). As shown in Figure S8, neither cocaine nor saline self-administration affected the number of trials omitted, regardless of sex or transgene status (all *Fs* < 2.102, all *ps* > 0.107). Cocaine self-administration reduced the latencies to collect the reward in females (session X TG: *F*_14,252_ = 2.965, *p* = 0.023; TG−: *t*_10_ = 6.540, *p* < 0.001; TG+: *t*_8_ = 4.122, *p* = 0.003) and males, regardless of transgene status (session: *F*_14,280_ = 4.607, *p* = 0.016; TG−: *t*_10_ = 3.215, *p* < 0.043; TG+: *t*_11_ = 4.191, *p* = 0.025). Animals self-administering saline underwent no changes in collect latencies (all *Fs* < 1.879, all *ps* > 0.198) (see Figure S9). Neither cocaine nor saline self-administration had any effect on choice latency (all *Fs* < 1.595, all *ps* > 0.068; see figure S10).

## Discussion

Here we report that chemogenetic inhibition of VTA dopamine neurons during acquisition and performance of the crGT prevents cue-induced risk taking in males, while enhancing it in females. The manipulation reduced motor impulsivity, but only in males. Despite these highly disparate effects across sex, this same manipulation protected both males and females against cocaine-induced deficits in decision making. Inhibition of VTA dopamine neurons also prevented cocaine-taking from reducing motor impulsivity.

Collectively, these results point to a complex interplay between reward-cues, dopaminergic signalling, risk taking, and the behavioural effects of cocaine. These results refute the notion that suppressing the dopamine system is universally beneficial or detrimental in terms of reducing addiction-like behaviour.

Before considering the potential significance of these findings, it is important to verify that the behavioural changes of interest are not the by-product of a general decrease in motor output, as may be expected from any manipulation that suppresses the dopamine system. During the crGT, TG+ve rats were slower to make a choice and collect reward, and also omitted more trials, indicative of reduced response vigour. However, only males showed a reduction in premature responses which accompanied a slight reduction in preference for the risky options over time, and these animals completed similar number of trials to their TG−ve counterparts. Female rats, on the other hand, showed no decline in premature responding and a significant increase in risky choice. It is therefore difficult to draw a link between indicators of motor slowing and decreases in motor impulsivity or risky decision-making patterns, as the former are clearly present in female rats without the latter.

In concordance with our initial hypothesis, we demonstrated a critical role for VTA dopamine neurons in cocaine-induced deficits in decision making (9). Specifically, we showed that self-administration of cocaine within the same diurnal period no longer resulted in an increase in risky choice when VTA dopamine neurons were inhibited during the crGT test session, even though rats were still self-administering ample cocaine. Following exposure to cocaine, recent data suggests dopamine neurons are rendered hypersensitive and more responsive to cues (31, 32). By chemogenetically inhibiting VTA dopamine neurons during task performance, we may have effectively dampened the ability of uncertain rewards and their accompanying audiovisual cues from driving the dopamine system, thereby preventing the descent into risky decision making.

We theorised that the repeated phasic dopamine bursts predicted to arise from daily streams of cued, uncertain rewards trigger an adaptive down-regulation of the dopamine system. We therefore predicted that chemogenetically inhibiting dopamine neurons system during each daily session should minimise the effect of the cues, and minimise risky decision making. While male rats did appear to adopt a slightly more optimal strategy, female rats instead dramatically increased preference for the risky options. The improved performance in males was also linked to the decrease in motor impulsive responses, such that the effect became non-significant when premature response rates are co-varied in the analysis. Chronically dampening the dopamine system in male rats may thusly reduce poor decisions mediated by impulsive “urges” but may not fundamentally improve outcome evaluation or action selection.

The marked increase in risky choice in females caused by chemogenetic inhibition of VTA dopamine projections, in the absence of any change in premature responding, was totally contrary to our predictions. However, a substantial body of evidence suggests that the mesocorticolimbic dopamine systems of females and males differ, both at baseline and in their respective responses to dopaminergic drugs (16, 18, 33). Numerous microdialysis experiments have demonstrated that, at baseline, males exhibit higher tonic levels of accumbal dopamine than females (34, 35). Taken together with the evidence that chemogenetic manipulations of VTA dopamine neurons produce similar changes in VTA activity and terminal dopamine release across the sexes (36), it is possible that the chemogenetic manipulation implemented here rendered absolute dopamine levels lower in females, as compared to males. Consistent with the inverted-U function which describes the dopaminergic contributions to some behaviours (37), inhibition of VTA dopamine neurons may have reduced dopamine to a level that promoted optimal performance in males, whereas in females, the manipulation may have reduced dopamine to detrimental levels (38). Indeed, extremely low levels of striatal dopamine are associated with impaired cognitive flexibility as well as the affective state of reward deficiency (39, 40). In females, cognitive inflexibility could have impaired learning of the task by causing rats to perseverate on risky options while failing to sample and switch to more optimal strategies. Furthermore, a state of reward deficiency may have increased the allure of the rewards with the highest payouts and most salient cues, as these options would presumably result in the most phasic dopamine release.

In rationalizing the sex differences we observed here, it is also important to consider that the VTA dopamine neurons we targeted project to numerous extrastriatal targets, including the prefrontal cortex (PFC) (41). Most complex cognitive behaviours, including decision making, are orchestrated in a large part by an interaction of dopaminergic processes between the PFC and striatum, and critically, the dopaminergic projection from the VTA to the PFC is denser in female rats than it is in males (42). Inhibiting all dopaminergic projections from the VTA may have therefore had a greater inhibitory effect on prefrontal dopaminergic processes in females, compared to males. Local antagonism of prefrontal D2 receptors causes significant deficits in probabilistic decision making (43). It is therefore possible that the decline in decision making ability we observed here in females were mediated by a greater reduction in prefrontal dopamine. Though estradiol powerfully influences dopamine neurotransmission in female rats (44), we do not suspect circulating gonadal hormones contributed to the sex differences we observe here. The longitudinal design of this study allowed for data collection over numerous estrous cycles, and as such, the data represent behaviour averaged over all stages of the cycle. We and others have furthermore shown that decision making in the crGT is not influenced by estrous stage (16, 17).

Chronic inhibition of VTA dopamine neurons increased cocaine intake, which matches the behavioural effect of administering low doses of dopamine antagonists (45). It is important to note that rats were no longer under the influence of CNO while self-administering cocaine, as the drug administration sessions took place over eight hours after injection of CNO. Nevertheless, this pattern of behaviour is consistent with enduring reductions in tonic dopamine levels. Indeed, others have demonstrated pharmacologically that chronic dopamine antagonism leads to long-lasting reductions in tonic extracellular dopamine levels (46, 47); these same neurobiological changes are thought to underlie the anhedonia associated with addiction in humans, with lower striatal *D*_2_ receptor levels associated with an increased propensity to self-administer cocaine (48, 49). Based on this evidence, the phenotype resulting from our present manipulation may have rendered animals relatively under-responsive to the dopaminergic stimulation produced by cocaine, leading animals to self-administer more drug in order to obtain desirable brain cocaine concentrations.

An alternative, and more speculative hypothesis concerns the potential synergy between cocaine self-administration and risky taking. We found previously that healthy animals trained on the crGT *either* self-administered more cocaine than animals trained on the uncued task, *or* made more risky decisions while self-administering cocaine in the same diurnal period (9). These data may suggest that making risky choices could somehow substitute for cocaine. Compared to control animals, rats that underwent inhibition of VTA dopamine neurons during daily crGT sessions experienced less on-task dopamine elevation. These rats may therefore consume more cocaine in an effort to boost accumbal dopamine levels, as sufficient stimulation may not have been achieved through the task.

Through combining daily cognitive behavioural test sessions with cocaine self-administration, the current study sought to evaluate how dampening dopamine signalling affected the synergistic interactions between drug-taking, impulsivity, and risky decision making in males and females. Although such an approach may seem unnecessarily complicated, the number of behaviours analysed still represents a small fragment of the cognitive processes that human drug users employ each day. By relying exclusively on simple behavioural models of addiction disorders, we risk drawing overly simplistic conclusions. The findings reported here strongly suggest that the ability of cocaine-taking to subsequently affect measures of both risky choice and impulsivity results from a disinhibition or hypersensitivity of the mesolimbic dopamine system; when we suppressed activity in this pathway, cocaine self-administration no longer altered these behaviours on the crGT. However, this same inhibition resulted in greater drug-taking, suggesting that any attempt to impede dopaminergic signalling in order to improve the cognitive sequelae of drug abuse may have unwanted consequences. Furthermore, although dampening dopaminergic activity during acquisition of the crGT had a generally positive impact on impulse control and decision making in male animals, this same manipulation drastically enhanced risky choice in females. Given that a rise in preference for uncertain options is a predictor of drug use (50), any putative therapeutics designed to prevent a “hyperdopaminergic state” from developing and contributing to the generation of addictions may be counter-productive, particularly in females. Despite decades of research into the relationship between dopamine signalling and addiction, questions clearly remain as to how activity within the dopamine system prevents or facilitates the addicted state. Combining complex behavioural models with sophisticated neural manipulations in animals will hopefully help to resolve these pressing issues, and lead to translationally meaningful results.

## ACKNOWLEDGEMENTS

The authors wish to thank Andrew Li for creating the LATEX pre-print and for editing this manuscript.

## AUTHOR CONTRIBUTIONS

TJH & CAW designed the experiment. TJH & KMH performed the surgeries. TJH, KMH, BAH, SAE, & CSC conducted the behavioural testing. CDH, CSC, & SK conducted the histology. CSC & TJH did the confocal imaging. GDB & BR provided statistical consultation. TJH & CAW wrote the paper.

