## Supplemental Figures for "Dopamine neurons gate the intersection of cocaine use, decision making, and impulsivity" (from another version of this preprint: version 1, 2020-06-30)

### Supplemental Figure 1. Hynes et al.

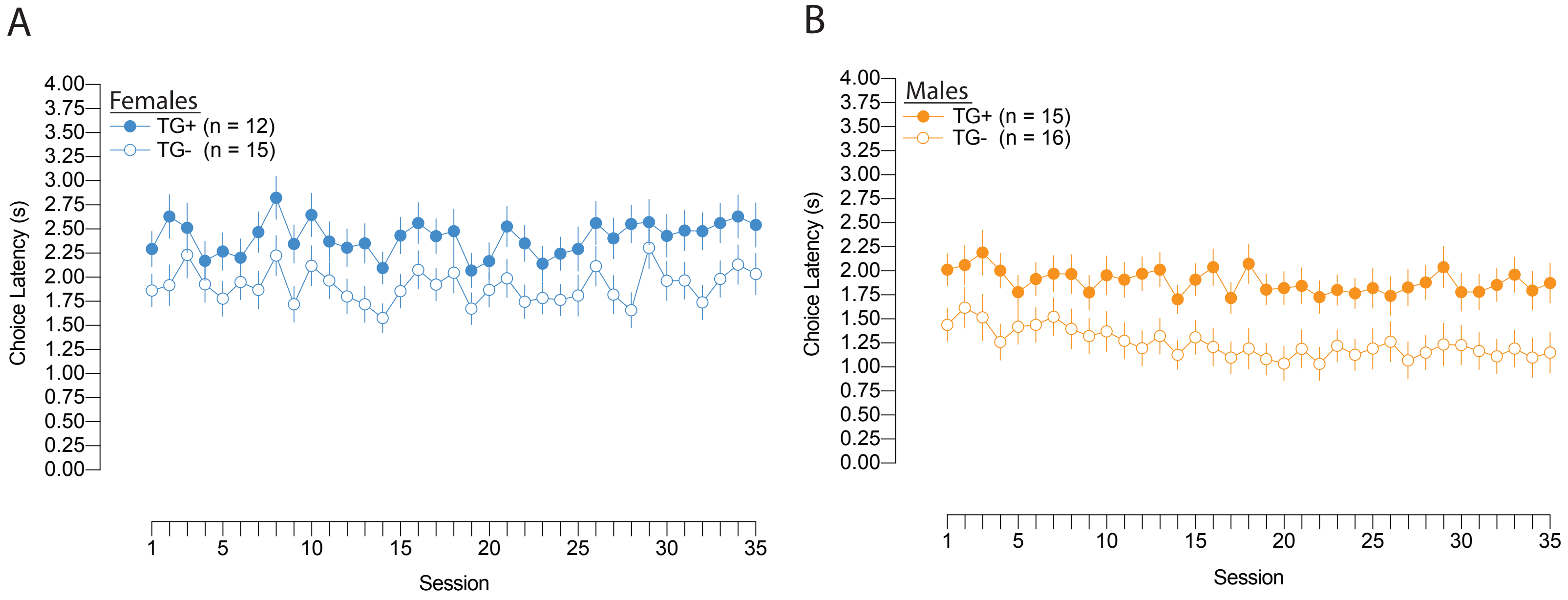


Figure S1. Longitudinal choice latencies during crGT training. In both (A) females and (B) males, TG+ rats were slower to make a choice. Regardless of transgene status, females were slower to make a choice than males. Data points represent group means +/- 1SEM.

### Supplemental Figure 2. Hynes et al.

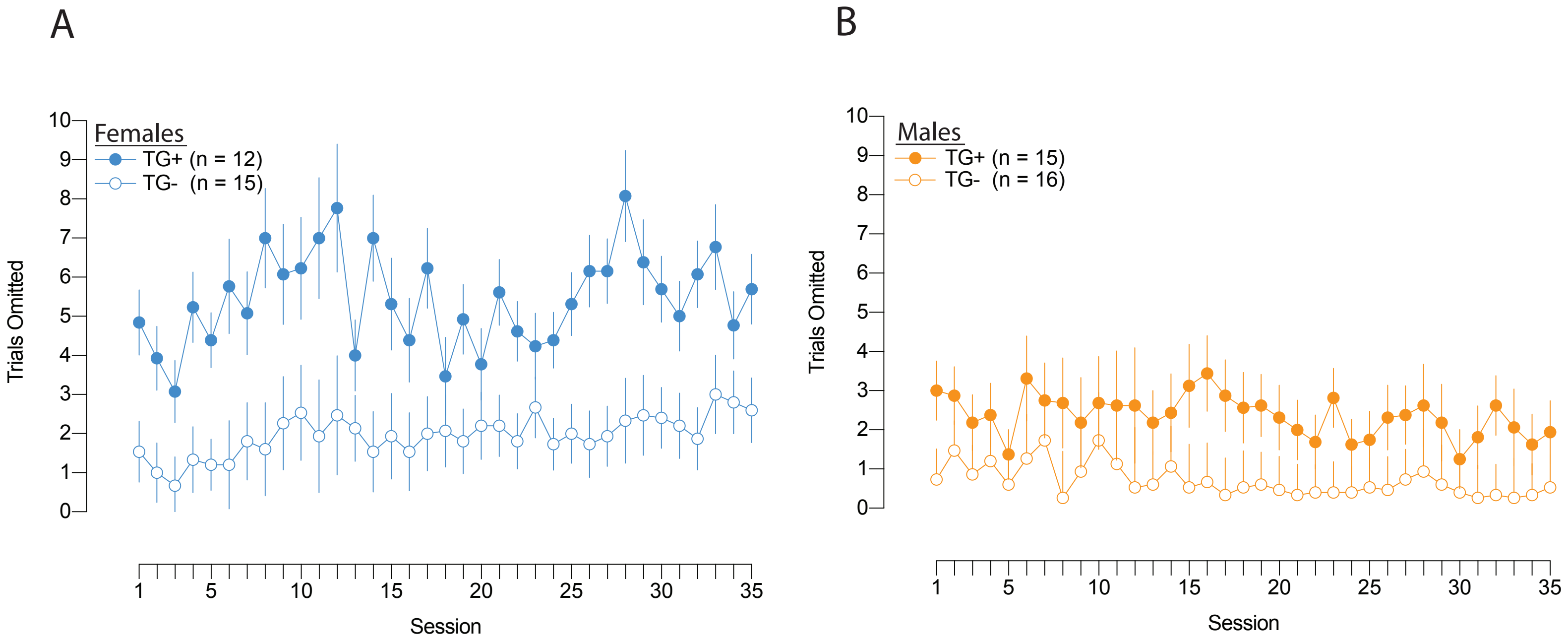


Figure S2. Longitudinal trial omissions during crGT training. In both (A) females and (B) males, TG+ rats omitted more trials than TG- rats. TG- males and females did not differ from one another. Data points represent group means +/- 1SEM.

### Supplemental Figure 3. Hynes et al.

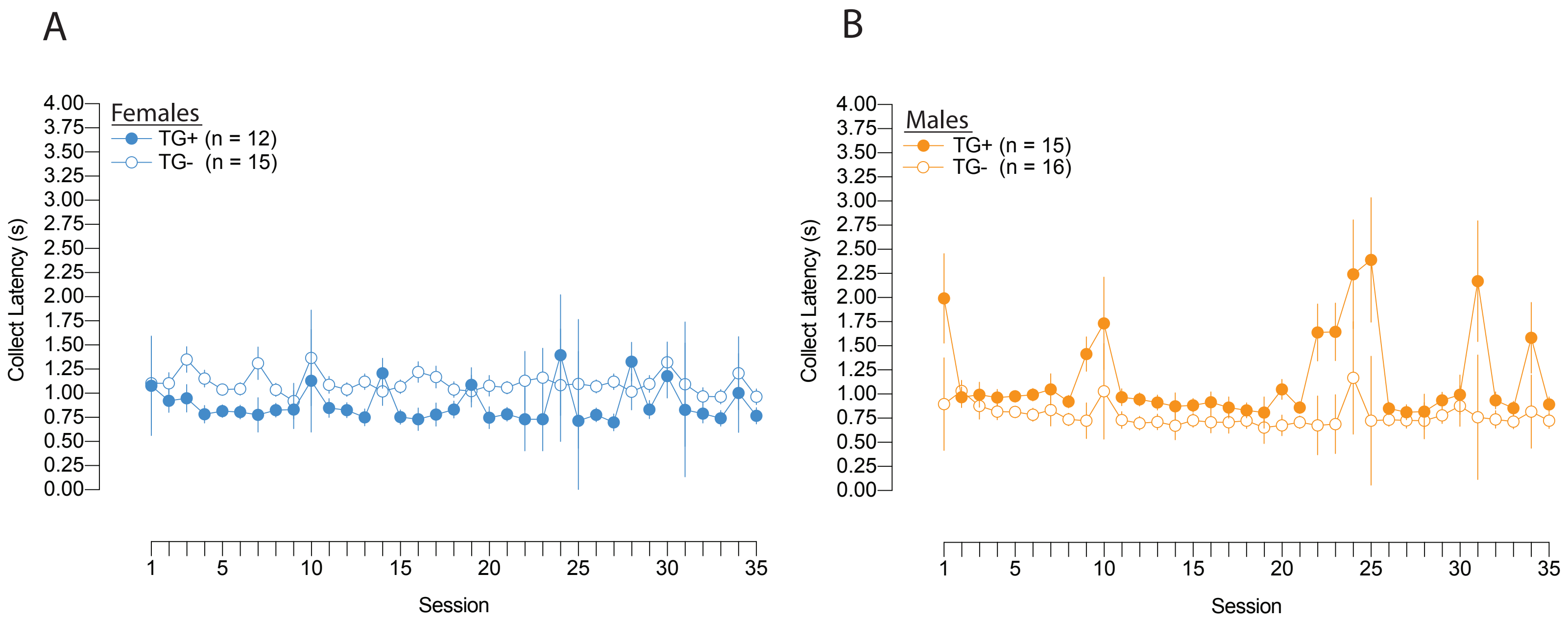


Figure S3. Longitudinal collect latencies during crGT training. (A) TG+ females were faster to retrieve the reward than TG- females, while (B) TG+ males were slower to do so than TG- males. TG- males and females did not differ from one another. Data points represent group means +/- 1SEM.

### Supplemental Figure 4. Hynes et al.

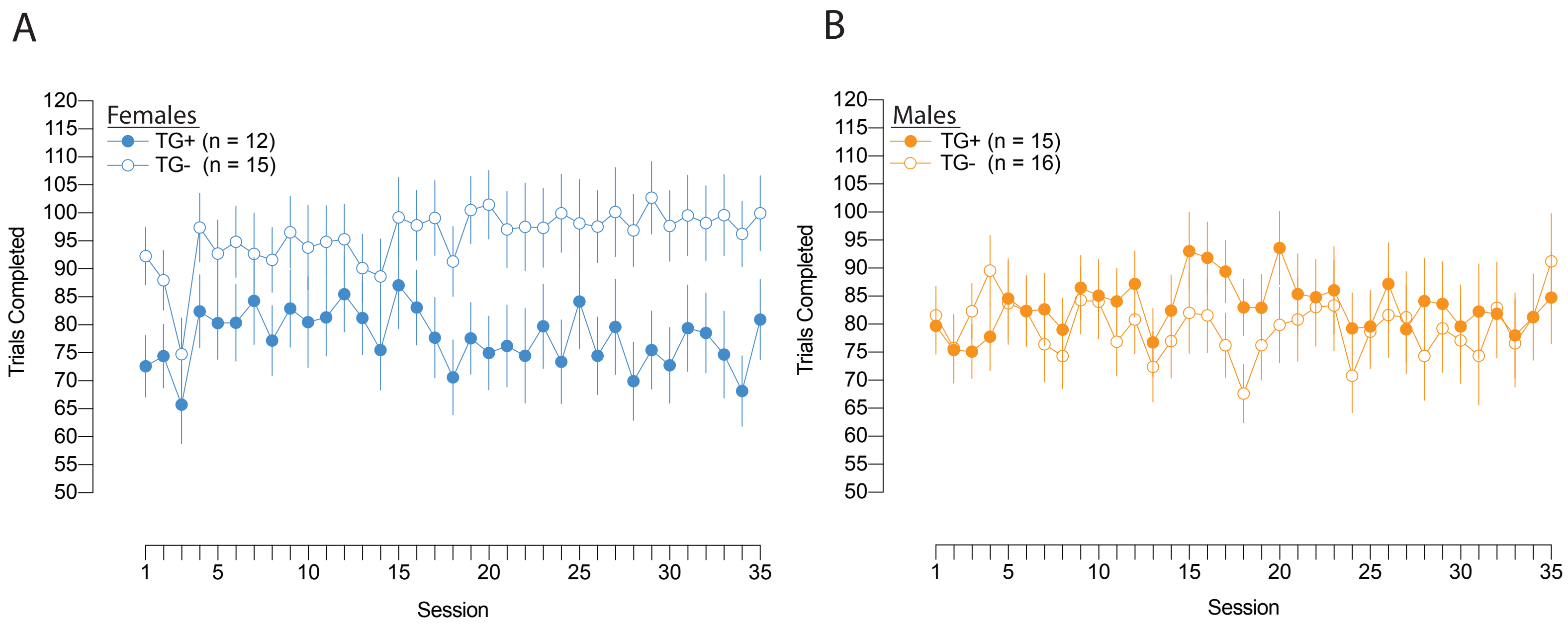


Figure S4. Longitudinal trials completed during crGT training. (A) TG+ females completed fewer trials per session than TG- females. (B) TG+ and TG- males did not differ from one another. Data points represent group means +/- 1SEM.
